# Long-access heroin self-administration induces region specific reduction of grey matter volume and microglia reactivity in the rat

**DOI:** 10.1101/2024.02.26.582024

**Authors:** Nazzareno Cannella, Stefano Tambalo, Veronica Lunerti, Giulia Scuppa, Luisa de Vivo, Sarah Abdulmalek, Analia Kinen, James Mackle, Brittany Kuhn, Leah C. Solberg Woods, Dongjun Chung, Peter Kalivas, Laura Soverchia, Massimo Ubaldi, Gary Hardiman, Angelo Bifone, Roberto Ciccocioppo

## Abstract

In opioid use disorder (OUD) patients, a decrease in brain grey matter volume (GMV) has been reported. It is unclear whether this is the consequence of prolonged exposure to opioids or is a predisposing causal factor in OUD development. To investigate this, we conducted a structural MRI longitudinal study in NIH Heterogeneous Stock rats exposed to heroin self-administration and age-matched naïve controls housed in the same controlled environment. Structural MRI scans were acquired before (MRI_1_) and after (MRI_2_) a prolonged period of long access heroin self-administration resulting in escalation of drug intake. Heroin intake resulted in reduced GMV in various cortical and sub-cortical brain regions. In drug-naïve controls no difference was found between MRI_1_ and MRI_2_. Notably, the degree of GMV reduction in the medial prefrontal cortex (mPFC) and the insula positively correlated with the amount of heroin consumed and the escalation of heroin use. In a preliminary gene expression analysis, we identified a number of transcripts linked to immune response and neuroinflammation. This prompted us to hypothesize a link between changes in microglia homeostasis and loss of GMV. For this reason, we analyzed the number and morphology of microglial cells in the mPFC and insula. The number of neurons and their morphology was also evaluated. The primary motor cortex, where no GMV change was observed, was used as negative control. We found no differences in the number of neurons and microglia cells following heroin. However, in the same regions where reduced GMV was detected, we observed a shift towards a rounder shape and size reduction in microglia, suggestive of their homeostatic change towards a reactive state. Altogether these findings suggest that escalation of heroin intake correlates with loss of GMV in specific brain regions and that this phenomenon is linked to changes in microglial morphology.

## INTRODUCTION

Opioid use disorder (OUD) is a chronic psychiatric condition characterized by compulsive drug taking, severe intoxications, and abstinence episodes generally followed by relapse. Over the last twenty years, the United States has witnessed a rise of overdose deaths caused by heroin and prescription opioids (NIDA, 2023) that have drawn renewed attention toward OUD. This called for research efforts to understand the neurobiology and neuropharmacology of OUD, necessary to widen the basis for future development of OUD treatment and safer opioid-based pain therapies.

Magnetic resonance imaging (MRI) studies in patients diagnosed with OUD consistently reported brain structural alterations compared to healthy controls. Grey matter volume (GMV) reduction in heroin dependent patients was observed in the prefrontal (Lin et al., 2018; Qiu et al., 2013; Shi et al., 2020), cingulate (Schmidt et al., 2021; Wang et al., 2012), and insular (Bach et al., 2019; Bach et al., 2021) cortices or in combinations of these areas (Liu et al., 2009; Sun et al., 2017; Sun et al., 2016). Fewer studies described GMV reduction in subcortical areas such as amygdala (Schmidt et al., 2021), globus pallidus (Shi et al., 2020), and putamen (Bach et al., 2021). In some cases, increased GMV in the somatosensory cortex (Shi et al., 2020; Sun et al., 2016) and caudate putamen (Schmidt et al., 2021) of these patients were also reported.

While there is a general agreement that patients with OUD exhibit a decrease in cortical grey matter volume (GMV), the specific regions involved, and the extent of the observed reduction differ across studies. The heterogeneity in the results reported by human studies could stem from differences in inclusion/exclusion criteria. For instance, the length of abstinence at the time MRI scans were acquired vary from no abstinence (Liu et al., 2009) to more than five drug free years (Shi et al., 2020; Wang et al., 2016). Some studies included patients under methadone/buprenorphine treatments (Bach et al., 2019; Liu et al., 2009), or included in medically assisted diacetylmorphine delivery programs (Schmidt et al., 2021). In other studies, GMV alterations were not found in the general population but could be observed only in groups filtered for genetic predisposition (Sun et al., 2017; Sun et al., 2016). The relative number of males and females is also a source of variability among studies, as women were often not included or underrepresented. Finally, there are several factors that can significantly impact GMV, some of which are difficult to predict, such as chemical impurities found in illicitly sold heroin, or factors not directly associated with the pharmacological effects of the drug, such as poor quality of life often associated with heroin use (Frischknecht et al., 2011; Puigdollers et al., 2004; Yen et al., 2011). In fact, it has been demonstrated that GMV in anterior cingulate cortex, medial prefrontal cortex, and insula, the three brain regions mainly affected by heroin, is positively correlated with physical and general health related quality of life (Hahm et al., 2019).

In summary, it is difficult to disentangle the effect of heroin exposure on GMV from that of environmental confounding factors inherent in human studies. Another important limitation in clinical imaging studies is the difficulty in determining whether the MRI changes observed in OUD patients are the consequence of prolonged drug exposure or are predisposing factors to the disease. One way to answer this question and to clarify the impact of environmental variables, is to conduct longitudinal studies with imaging data collected prior and after opioid exposure and dependence development. For obvious reasons, conducting these studies in humans can be challenging, while it is relatively straightforward to carry them out in laboratory animals.

In the first part of our study, we used a longitudinal approach and structural MRI to measure GMV in rats before and after protracted (12 hrs. a day) exposure to operant heroin self-administration. Age matched heroin naive rats were used as a control. The study was carried out under tightly controlled environmental conditions. To better mimic the genetic heterogeneity of the human population, we used NIH heterogeneous stock (HS) rats, an outbred line classically used for genetic and phenotypic studies including GWAS analysis (Solberg Woods and Palmer, 2019). Previous research has shown that these rats are suitable for studying opioid abuse-related behavior (Allen et al., 2021; de Guglielmo et al., 2019; Kallupi et al., 2020; Kuhn et al., 2022). Behavioral measures aimed at evaluating the motivation for heroin and the transition from its use to abuse were also taken. In the second part of the study, guided by the MRI results and transcriptomic data from HS rats exposed to heroin, which suggested neuroimmune system activation, we conducted an extensive immunohistochemistry investigation to assess whether changes in GMV following heroin use involved alterations in microglia morphology compared to neurons.

## MATERIALS AND METHODS

### Animals

NIH heterogeneous stock (HS) rats were obtained from the Biomedical Resource Center, Medical College of Wisconsin (Milwaukee, USA). Rats were housed in groups of two, in a room with artificial 12/12 h light/dark cycle (light off at 8 AM), constant temperature (20-22°C) and humidity (45-55%). Rats had *ad libitum* access to tap water and food pellets (4RF18, Mucedola, Settimo Milanese, Italy) unless differently indicated. All procedures were conducted in adherence to the “European Community Council Directive” and “NIH Guidelines” for Care and Use of Laboratory Animals.

### Drugs

Heroin (diacetylmorphine hydrochloride, NIDA drug supply program) was dissolved in sterile physiological saline.

#### Experiment 1: Heroin self-administration and seeking

Surgeries, self-administration apparatus and food-reinforced pre-training of operant lever responding are described in **Supplementary Information.** Fifteen HS rats (7 males and 8 female) were implanted with an indwelling catheter into the right jugular vein to receive heroin infusions. Operant heroin self-administration (HSA) was carried out according to a protocol adapted from (Vendruscolo et al., 2011) and briefly described below. Additional ten rats (5 males and 5 female) were subjected to the occlusion of the right jugular vein and received i.v. saline infusions. This yoked group served as a control for the MRI experiment described below. Timelines of the experiment are depicted in **Figure 1A**.

**Figure 1.**
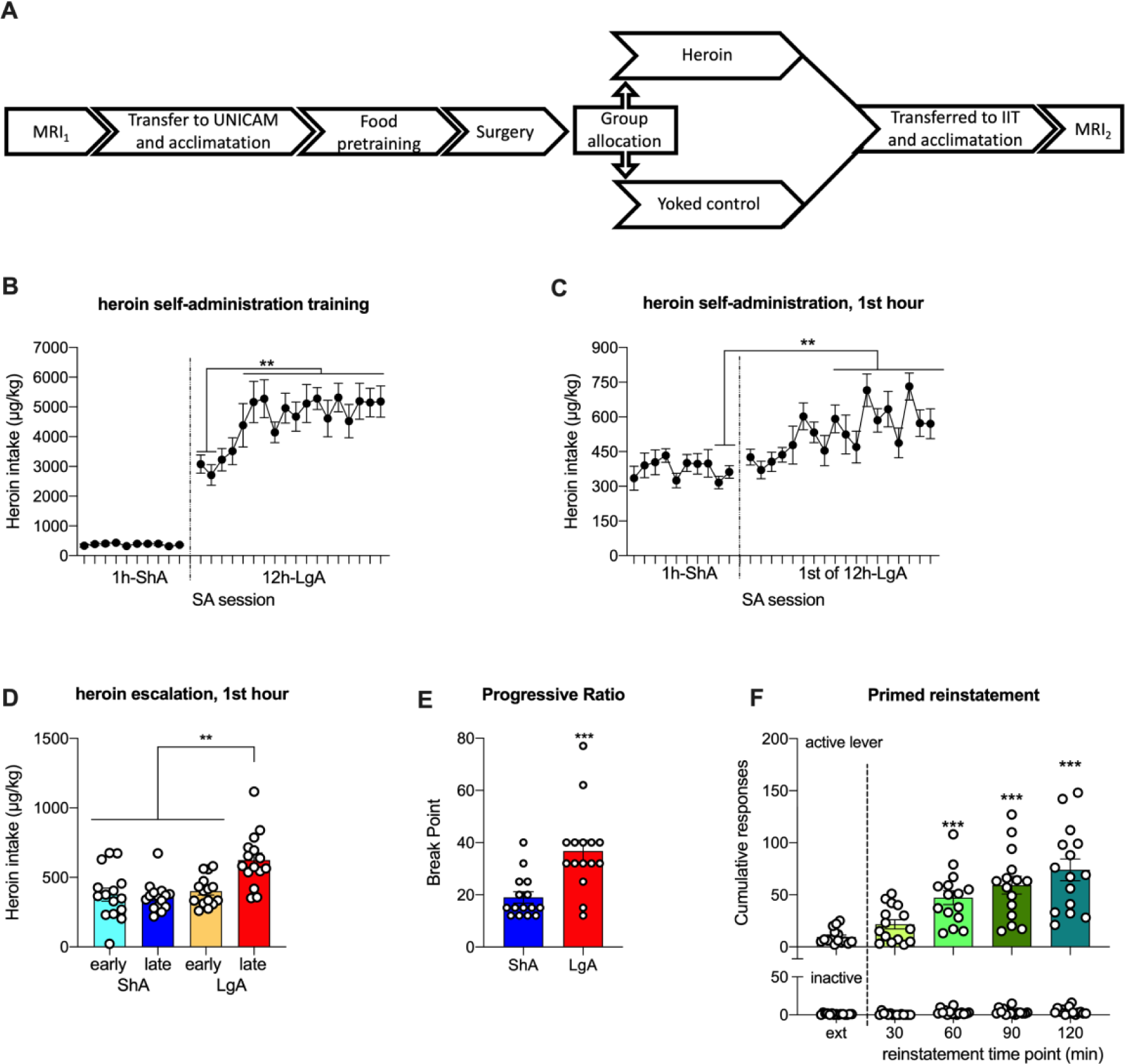
**A**) Flowchart of the longitudinal MRI experiment timeline. **B**) Heroin intake during ShA and LgA self-administration training. Rats showed stable heroin intake during the 1h ShA phase (left side of dashed line). During the 12h LgA phase heroin intake escalated over time, becoming statistically higher than the first two LgA session starting from the 6^th^ session, when it reached a plateau. **C**) Heroin intake during the 1h ShA sessions and the 1^st^ hour of the 12h LgA sessions. During the LgA phase the 1^st^ hour intake increased over time becoming statistically different from the last two days of ShA from the 6^th^ session. **D**) Escalation of heroin intake expressed as the difference between the average intake at four key time points: first three and last three ShA sessions (early and late ShA respectively), the first three and last three LgA sessions (early LgA and late LgA respectively). The intake in late LgA was significantly higher than the other three time points. **E**) Motivation for heroin expressed by the break point reached in PR sessions run during the ShA and LgA phase. Rats showed increased break point after LgA training. **F**) Heroin primed reinstatement of heroin seeking in a within session extinction-reinstatement protocol. Heroin priming reinstated active lever pressing, which increased over time compared to the last 30 minutes of extinction (upper panel). Inactive lever remained constantly low and was not statistically different from extinction (lower panel). Data are expressed as mean ± SEM. Significant differences: A-C) **p<0.01; D) ***p<0.001 vs ShA; E) *** p<0.001 vs extinction (ext), °p<0.05, °°p<0.01 and °°°p<0.001 vs previous reinstatement time point.

The heroin group was initially trained to 1-hour short access (ShA) HSA. Sessions started with the insertion of both retractable levers. The right lever was designated as the active lever. Depression of the active lever under fixed ratio 1 (FR1) schedule of reinforcement activated the infusion pump for 5 seconds. Each pump activation resulted in the intravenous delivery of 60 µg/kg of heroin in a volume of 0.1 ml. A 20 second time out (TO) followed pump activation. During TO, active lever pressing had no scheduled consequences. The cue light located above the active lever was activated contingently with the infusion pump and remained on for the whole TO. The left lever was designated as the inactive lever and functioned as control for non-specific behavior. Responses at the inactive lever were recorded but had no scheduled consequences.

After nine FR1 ShA SA sessions, motivation for heroin was evaluated under progressive ratio (PR) schedule of reinforcement similarly to that we previously described (de Guglielmo et al., 2015). In this session the response requirements necessary to receive a single reinforcement increased after every infusion according to the equation: [5e^(injection^ ^numbers^ ^x^ ^0.2)^]-5. The last ratio completed was defined as the break-point (BP) and was taken as a measure of the motivation for heroin expressed by the rat (de Guglielmo et al., 2015; Richardson and Roberts, 1996). The session ended after four hours in total had elapsed or if one hour passed since the last infusion earned, whichever occurred first.

After the PR test, rats were subjected to one additional ShA session, after which they entered the 12-hours long access (LgA) HSA phase. LgA sessions were identical to ShA sessions except that they lasted for twelve hours. <u>A</u>fter eight LgA sessions, rats were tested again in a PR schedule of reinforcement as described above for the ShA phase.

The day after the second PR test a heroin primed reinstatement test was run according to a within session extinction-reinstatement protocol (Shaham et al., 2003). This session lasted three hours in total. During the first hour, the syringe pumps were off and lever pressing did not deliver heroin. This session served as an extinction phase characterized by progressive decrease of lever pressing. At the beginning of the second hour, syringe pumps were switched on for five minutes, during which rats could self-administer a maximum of two infusions in the same condition of a SA session. If a rat failed to self-administer two heroin infusions within five minutes, the infusions were delivered by the operator that manually activated the infusion pump. The two infusions served to prime heroin seeking behavior that was monitored for the remaining 2 hrs. After the priming reinstatement test, rats returned to LgA FR1 baseline self-administration for an additional ten sessions. Heroin training and behavioral tests, sessions were run for four days a week, from Monday to Friday with a random one-day break in within. Sessions were performed during the dark phase of the light/dark cycle, and during long access training tap water and chow-pellets were available in the self-administration chambers. The yoked group received 35 random saline infusions paired with the same cues experienced by the heroin rats.

#### Experiment 2: Effect of heroin consumption on grey matter volume in HS rats

<u>At</u> fourteen-week of age, prior to enter into Experiment 1, twenty five HS rats (13 males and 12 females) were subjected to the first MRI acquisition (MRI_1_) at the Italian Institute of Technology (IIT, Rovereto). A week later, rats were transferred to the University of Camerino. At twenty weeks of age rats were divided into the heroin group (7 males and 8 female) and yoked saline controls (5 males and 5 female) and subjected to Experiment 1. At completion of Experiment 1 rats were transferred back to the IIT for a new MRI acquisition (MRI_2_). MRI_2_ acquisition started at 5 days and ended 13 days after the last heroin self-administration (SA) session. Experimental timeline is summarized in **Figure 1A**. For MRI acquisition a 7 Tesla (T) Bruker Pharmascan (Bruker BioSpin, DE) in a double coil configuration was used. A 72-mm i.d. single channel transmission-only resonator was actively decoupled with a four-channel phased-array receive-only surface coil optimized for the rat brain. After a scout image, high resolution T2w RARE images (TR = 5500 ms, TE = 76 ms, NEX = 8, MTX = 256 × 256 × 25, FOV = 35 × 35 × 25 mm, ACQtime = 7min40s) were acquired for voxel-based morphometry analysis. T2w RARE was used with a multislice encoding (2D). Accurate and consistent positioning of the FOV was ensured by placing the center of the first slice 0.5mm rostral to the rhinal fissure. This allowed to cover the entire forebrain and midbrain regions.

#### Experiment 3: Effect of heroin consumption on the morphology of microglia in HS rats

Different groups of HS rats (3 males and 3 females) trained to LgA heroin self-administration and saline-yoked controls (2 males and 2 females) were used (**Supplementary Figure S1**) for immunohistochemistry experiments. Specifically, nine days after the last SA session, corresponding to the average time point at which the MRI_2_ of the longitudinal MRI experiment was acquired, the heroin and saline exposed rats were deeply anesthetized with 5% isoflurane and perfused transcardially with a flush (∼30 s) of saline followed by 200 ml of 4% paraformaldehyde (PFA) in phosphate buffer (PB) and post-fixed overnight in PFA at 4°C. Brains were subsequently sectioned with a vibratome (Leica) into 50 μm thick coronal slices, collected in PB and 0.02% sodium azide (NaN), and stored at 4°C.

Four to five slices for each selected region were collected: the medial prefrontal cortex (mPFC) centered at bregma (+2.20), that included the Cingulate area 1 and the Prelimbic Cortex (Cg1, PrL); the Primary Motor Cortex (M1), centered at bregma (+2.20); the Insular cortex (IC), centered at bregma (+0.70), that included the Granular, Dysgranular and Agranular Insular Cortex (GI, DI, AID, AIV) (**Supplementary Figure S2**). Coordinates refer to Paxinos & Watson (1998) (Paxinos and Watson, 1998).

Neuronal nuclei and parvalbumin positive neurons were double stained with NeuN and anti-parvalbumin (PV) antibodies respectively; Iba1 antibody was used instead as a marker of microglia.

For NeuN and PV double-staining (10 animals, 2 sections/animal), sections were washed three times with phosphate-buffered saline (PBS, pH 7.4), treated with a blocking solution of 3% normal goat serum (NGS) and 0.1% Triton X-100 (TXT-100) for 1h at room temperature (RT), and then incubated in the blocking solution containing mouse anti-NeuN antibody (1:500, Synaptic Systems 266 004) and guinea pig anti-parvalbumin antibody (1:1000, ImmunoStar 24428) for 2h at RT and then overnight at 4°C. The day after, sections were washed three times with PBS and treated with a blocking solution (3% NGS) for 20 min at RT and incubated with 3% NGS PBS containing secondary antibodies (1:600, Alexa-Fluor 594 anti-mouse IgG, Invitrogen and Alexa-Fluor 488 anti-guinea pig IgG, Invitrogen) for 90 min at RT. For Iba-1 staining (10 animals, 2 sections/animal), sections were washed three times with PBS (pH 7.4), treated with a solution of 3% bovine serum albumin (BSA) and 0.3% Triton X-100 for 1h at RT, and then incubated in the blocking solution (1%BSA and 0.1% TXT-100) containing rabbit anti-Iba1 primary antibody (1:600, Fujifilm Wako 019-19741) for 2h at RT and overnight at 4°C. Sections were then washed three times with PBS and treated with a blocking solution (1% BSA) for 20 min at RT and then, they were exposed for 90 min to 1% BSA in PBS containing a secondary antibody (Alexa Fluor 488 anti-rabbit IgG, Invitrogen) at RT.

Finally, all sections were washed with PBS, mounted, air-dried and cover slipped using either Vectashield mounting medium (NeuN and PV) or Vectashield mounting medium with DAPI (Iba1). In control slices primary antibodies were omitted.

Sections were examined with a confocal microscope (Nikon C2+Laser Scanning confocal) and acquired as 512x512 pixel images (0.41 µm pixel size; Z-step, 1 µm; 210.12x210.12 microns) using a 60x oil immersion lens for Neun/PV sections, and an air-dry 40x lens for Iba1 sections. Six fields/layer/animal in layers III and V were acquired for each region (mPFC, M1, IC).

Images for NeuN and PV were analyzed using FIJI ImageJ software (Bellesi et al., 2017). The number of NeuN and PV positive cells was counted manually. PV morphology was assessed by brain area around each PV positive cell and measuring perimeter, area, and density. Image processing of Iba-1 was performed using a custom-made script to measure individual microglial area and perimeter length in MATLAB, as described in (Bellesi et al., 2017).

### Data analyses

#### Behavioral Experiments

Food and heroin self-administration, escalation of heroin intake, and primed reinstatement were analyzed by one-way ANOVA with time as repeated measure. Break points reached during the two PR sessions were compared by Wilcoxon match paired signed rank test. Statistical significance was conventionally set at p<0.05. Data are presented as mean ± SEM.

#### Voxel Based Morphometry

T2w RARE images, acquired at high resolution, were analyzed to investigate voxelwise differences in local GMV using a modified version of the FSL-VBM tool of FSL (Smith et al., 2004) to account for rat brain specific issues (Tambalo et al., 2015). Briefly: an unbiased GM template was created by averaging the entire group at both time points (experimental: n = 30, control: n=20). Individual GMV were registered to the unbiased template using the FSL FNIRT algorithm (Schnabel et al., 2003) and then modulated to correct for local deformations by the Jacobian of the warp field, as reported in (Ashburner and Friston, 2000). The modulated GM images were then smoothed with an isotropic Gaussian kernel (σ = 3 mm), and a nonparametric permutation test was applied. The null distribution for the data in the VBM statistics was built over 5,000 permutations. To define more precisely the distribution of GM density modulation in different anatomical regions of the brain, a volumetric reconstruction of the digital anatomical rat brain atlas (Paxinos and Watson, 1998) was coregistered with GM images in template space.

#### Histochemical Analyses

Cell density of NeuN, PV and Iba1 positive cells were compared between heroin experienced and control rats using the non-parametric Mann-Whitney test. Cumulative distributions of morphological parameters of Iba1 and PV positive cells were compared between heroin experienced and control rats using the non-parametric Kolmogorov-Smirnov (KS) test. Male and female rats were grouped together for analyses.

## RESULTS

### Experiment 1: Heroin self-administration and seeking in HS rats

<u>After</u> food-reinforced pretraining (**Supplementary Figure S3**), rats rapidly acquired ShA HSA responding [F(9, 14)=1.0; p>0.05] (**Figure 1B**, left to the dashed line). Responding at the inactive lever was always very low and did not change during training [F(9, 14)=0.9; p>0.05]. Then, when rats entered the 12-hour long access (LgA) phase, ANOVA revealed an overall effect of SA sessions [F(17, 14)=5.9; p<0.001], with heroin intake increasing over time. The increase in intake became statistically significant starting from the 6^th^ LgA session, after which heroin intake remained stable (**Figure 1B**, right to the dashed line). Responding at the inactive lever remained stable over time [F(17, 14)=1.1; p>0.05]. This indicates an escalation of heroin intake induced by long access to heroin, a marker of drug dependence (Ahmed and Koob, 1998; Vendruscolo et al., 2011). Escalation is also indicated by an increase in intake during the first hour of the LgA sessions (Ahmed and Koob, 1998), therefore we analyzed the heroin intake at this time point comparing heroin intake under ShA and LgA condition and we found an overall effect of sessions [F(27, 14)=6.0; p<0.0001]. Heroin intake at the first hour was stable during the ShA and the first four days of LgA, and it started to increase from the fifth LgA session. During the last seven LgA sessions, heroin intake was significantly higher than the last two days of ShA (**Figure 1C**). Analysis of inactive lever responding found no overall effect of sessions [F(27, 14)=1.0; p>0.05], indicating that the increase observed in the heroin intake did not derive from a general change in behavior and can be interpreted as an escalation of heroin intake. To further evaluate the magnitude of escalation, we compared the average heroin intake during the first hour of SA at four key time points: the first three ShA sessions (early ShA), the last three ShA sessions (late ShA), the first three LgA sessions (early LgA), the last three LgA sessions (late LgA). The ANOVA found an overall effect of time point [F(3, 14)=11.2; p<0.001]. Bonferroni’s post-hoc analysis revealed that the intake at late LgA was significantly higher than the other three time points (**Figure 1D**).

#### Motivation for heroin under PR measured after ShA and LgA HSA

Motivation for heroin was evaluated by analyzing the break points (BP) reached in the two PR SA sessions run at the end of the ShA and LgA sessions, respectively. A two-tailed Wilcoxon matched-pairs signed rank test revealed a significant difference between the two BP [W(15) = 102.0; p<0.001] (**Figure 1E**), indicating that rats showed a higher motivation for heroin after escalation of intake.

#### Heroin primed reinstatement

Finally, a within session extinction-priming reinstatement test was run. To analyze the reinstatement, data were computed in 30 min bins to compare the cumulative number of lever presses after priming with lever pressing occurring during the second half of the extinction phase. An ANOVA of active lever presses found an overall effect of time [F(4, 14) = 38.03; p<0.0001]. The number of presses increased over time during the reinstatement phase and Bonferroni’s post-hoc analysis revealed that it became significantly higher than extinction at 60 minutes into the reinstatement phase (p<0.001). Bonferroni’s test further revealed that at each time point within the reinstatement phase the number of presses was higher than the previous time point (**Figure 1F upper panel**). Analysis of the inactive lever also found an overall effect of time [F(4, 14) = 8.8; p<0.01]. Bonferroni’s post-hoc analysis revealed significant differences exclusively within the reinstatement phase time points, specifically between 120 and 60 minutes and between 120 and 30 minutes (**Figure 1F** lower panel). Altogether, these analyses indicated a significant priming-induced reinstatement of heroin seeking.

### Experiment 2: Effect of heroin consumption on grey matter volume in HS rats

In heroin experienced rats we observed a negative difference between GMV in MRI_2_ and MRI_1_ indicating a decrease in GMV in several cortical and subcortical regions. Specifically, GMV reduction was detected in the insular cortex (IC), prelimbic cortex (PL), infralimbic (IL), and Cingulate area 1 (Cg1), the caudal portion of the Retrosplenial Cortex (RsC), Orbitofrontal Cortex (OFC), Hypothalamus (Hyp), Thalamus (Th), Entorhinal Cortex (EnC), Hippocampus (Hipp), bed nucleus of the stria terminalis (BNST), Amygdala (amy), Nucleus Accumbens (Acb), and Dorsal Striatum (DS). (**Figure 2**, corrected- p < 0.01, blue = reduction of GM volume). No positive difference between GMV in MRI_2_ and MRI_1_ (i.e., no increase in GMV) was observed at any brain level. The heroin naive group of animals, evaluated at the same timepoints, showed neither positive nor negative GMV changes between the two scans, indicating that GMV was not affected by manipulation of the rats and/or aging When we used drug seeking and taking as covariates to assess their correlation with GMV changes (MRI_2_ minus MRI_1_) (**Figure 3**) we found a negative correlation with both heroin intake and escalation (both at the 1st and 12th hour of HSA) in the mPFC, IL, IC, DS and Acb. For IC and DS, this correlation extended also to the extinction responding and priming induced reinstatement of heroin seeking. Finally, in the caudal RsC GMV negatively correlated with priming and total heroin intake at the 12th hour.

**Figure 2.**
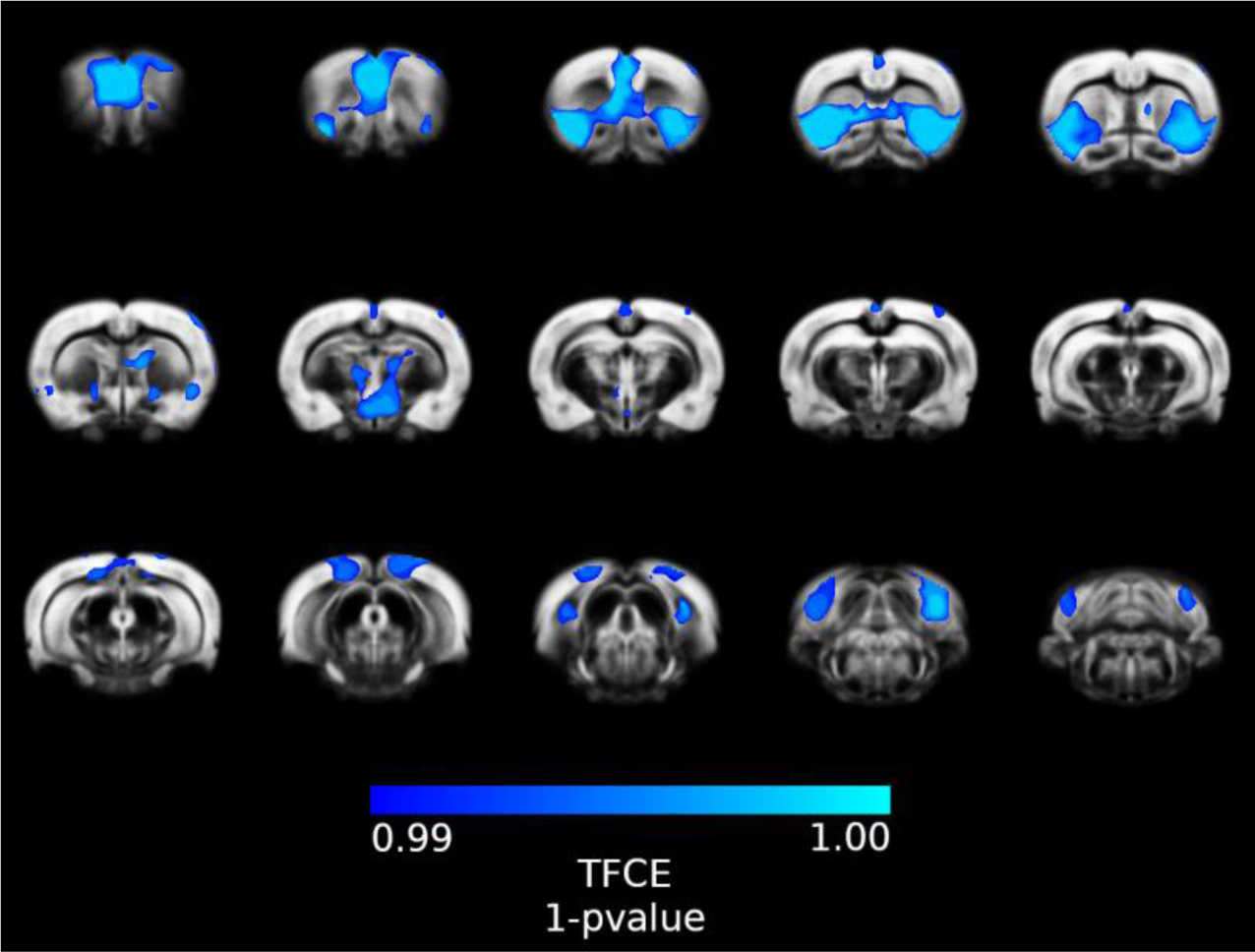
Spatial distribution of negative modulation of grey matter volume after drug exposure. Significant reduction of grey matter volume is thresholded at 1 - p_value_ > 0.99, TFCE corrected.

**Figure 3:**
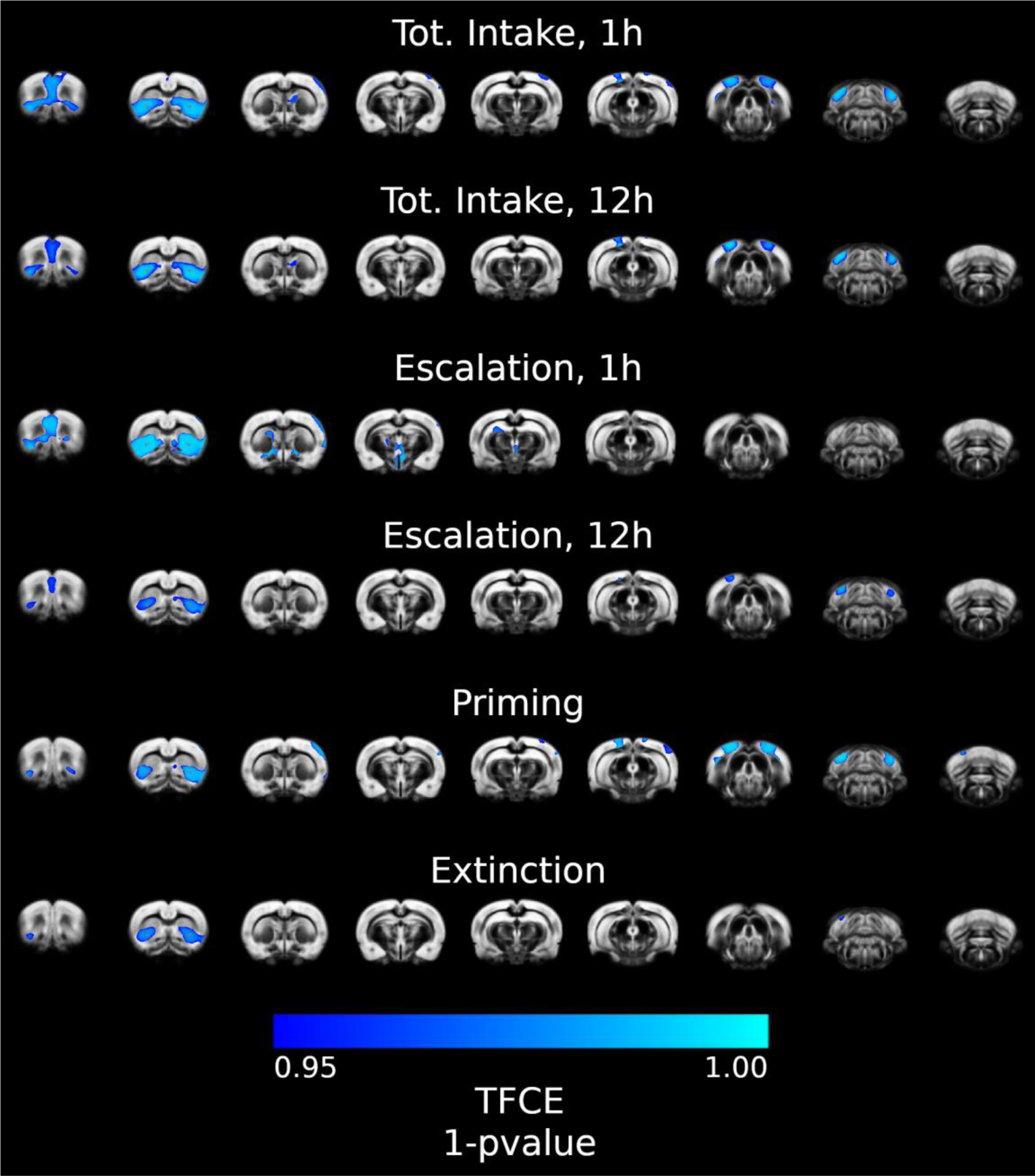
Voxelwise correlation maps of drug seeking and taking behavioral readouts against grey matter volume MRI2 - MRI1 difference. Significant negative correlations are thresholded at 1 - pvalue > 0.95, TFCE corrected.

### Experiment 3: Effect of heroin consumption on the morphology of microglia in HS rats

To further investigate the biological phenomenon that may be at the basis of the reduction in GMV observed in heroin experienced rats relative to controls, we measured the density and size of neurons and microglial cells in two brain regions in which we observed GMV reduction - the mPFC and the IC - and one region in which GMV reduction was not observed, the primary motor cortex (M1) (**Figure 4 A-C**).

**Figure 4.**
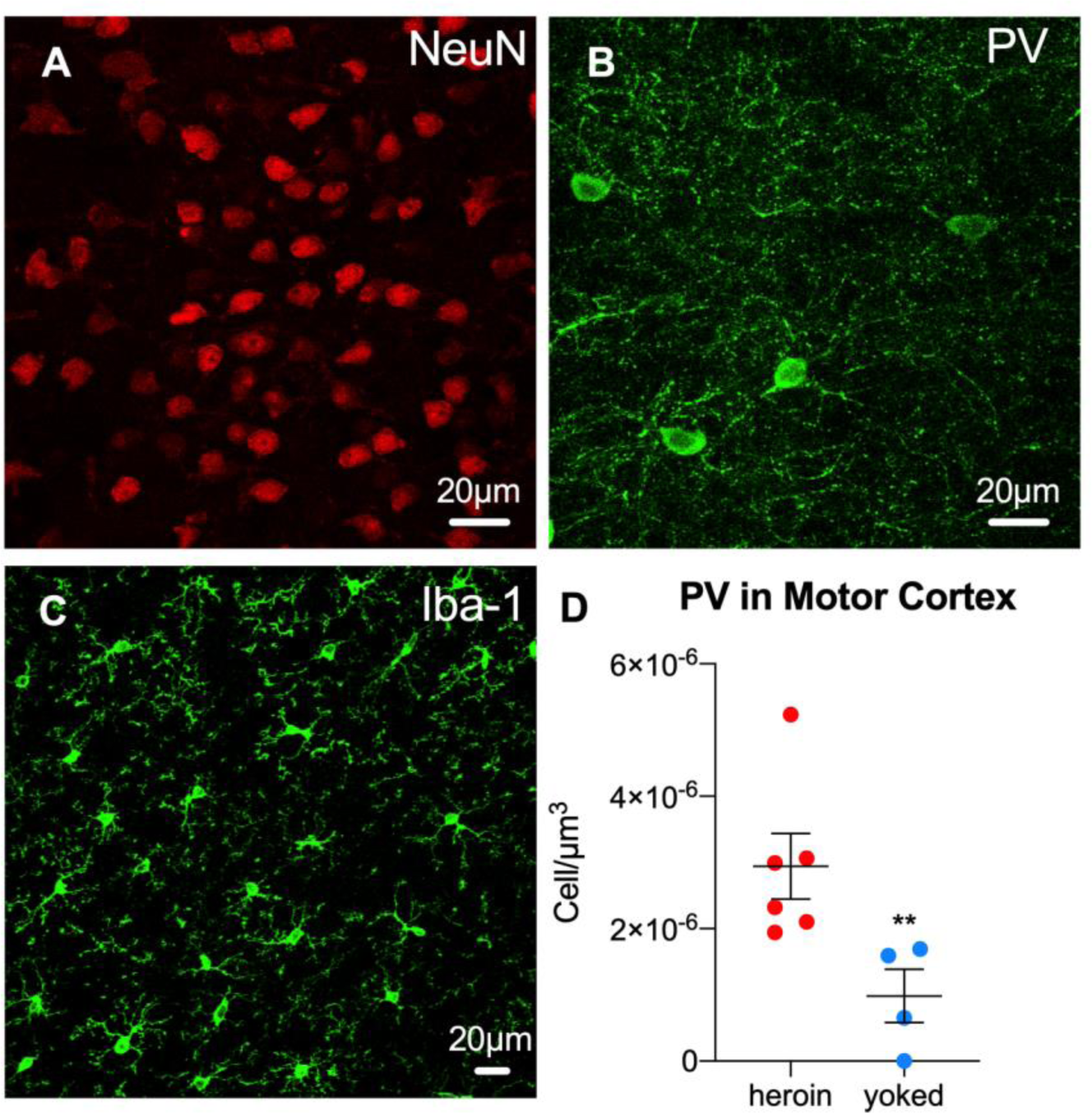
Representative image of **A**) NeuN staining and **B**) PV staining taken in the Insular Cortex and medial Prefrontal Cortex of a heroin treated rat, respectively; **C**) Iba1 staining taken in the Insular Cortex of a control rat. **D**) Heroin experienced rats showed higher PV-positive cell density in the Primary Motor cortex. **A-C**: scale bars = 20 µm; **D:** whiskers represent mean ± SEM, statistical significance **p < 0.01.

#### Cell density

Heroin experienced rats showed higher density of PV positive neurons in the M1 [Mann-Whitney U = 0; p< 0.01] (**Figure 4 D**), while no significant differences were found between heroin and saline yoked groups in the other brain regions and for the other cell types (**Supplementary Figure S4**).

#### Size of parvalbumin neurons

We estimated the area and perimeter of 701 PV positive neurons (214 mPFC, 223 IC, 264 M1) in heroin-exposed animals and 232 PV positive neurons (93 mPFC, 96 IC, 43 M1) in yoked controls. In the insula, we observed a rightward shift of both Area [KS D = 0.33; p<0.0001, +30.3%] and Perimeter [KS D = 0.27; p<0.0001, +12.9%] distributions in heroin-exposed animals relative to controls, indicating an increase in the size of these neurons (**Supplementary Figures S5**). Conversely, no significant change was observed in PV size in the mPFC and M1.

#### Morphological analysis of microglia

Here, we quantified cell size, perimeter length, and the area/perimeter ratio of 1236 Iba1 positive cells (375 mPFC, 452 IC, 409 M1) in heroin exposed animals and 1044 cells (327 mPFC, 357 IC, 360 M1) in yoked rats as a proxy of microglia state in three different brain regions. In the mPFC, we observed a reduction in the area [KS D = 0.39; p<0.0001] and perimeter length [KS D = 0.39; p<0.0001] of microglial cells, as shown by the leftward shift of both cumulative distributions, in the heroin experienced group relative to the saline yoked rats (**Figure 5A, B**). The area/perimeter ratio was instead right-shifted in the heroin experienced rats [KS D = 0.29; p<0.0001] (**Figure 5C**). The higher area/perimeter ratio observed in heroin experienced rats, in which both area and perimeter were smaller, is consistent with a larger decrease observed in cell perimeter (−58.6%) than in cell area (−50.5%).

**Figure 5:**
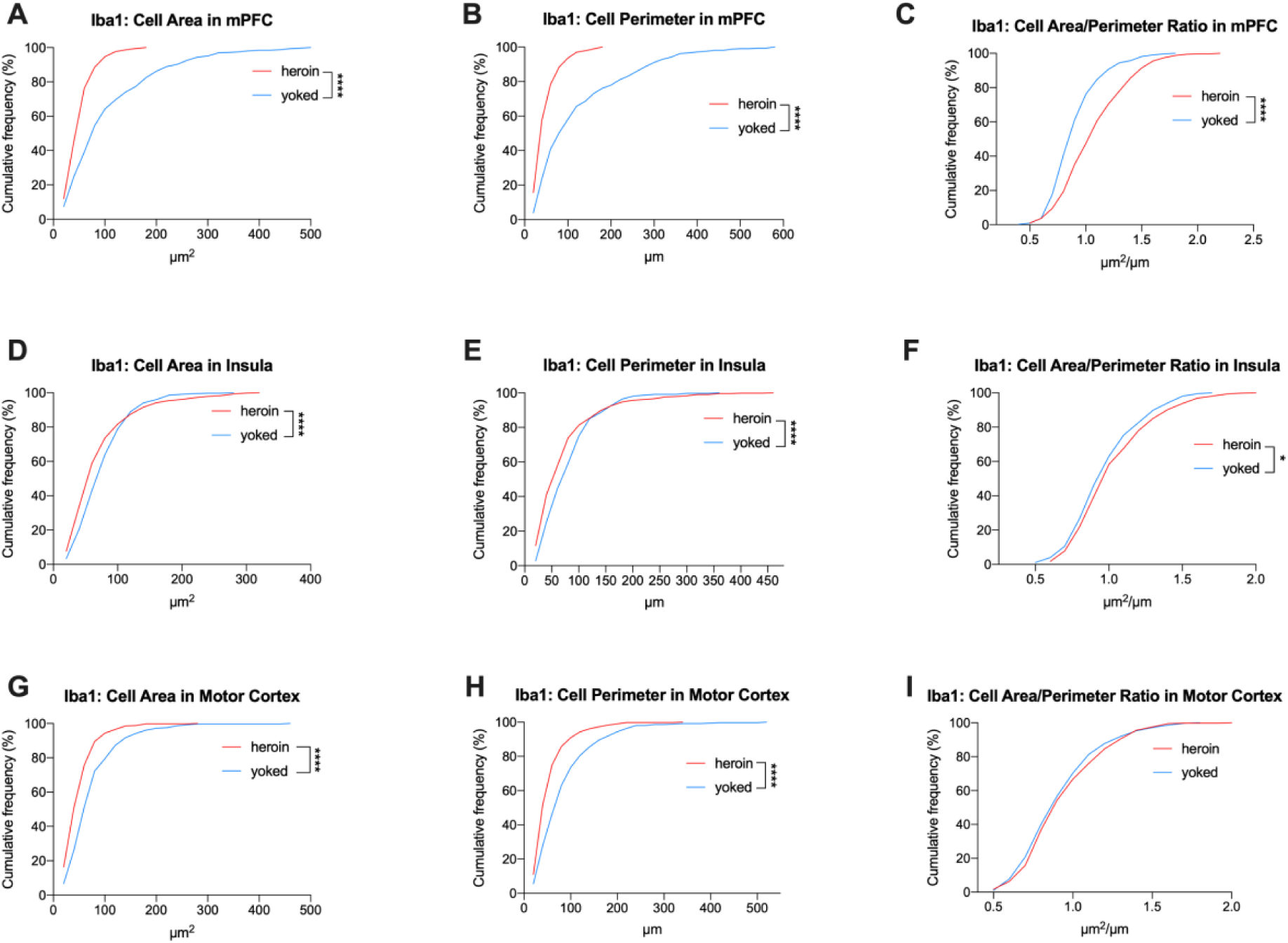
Cumulative distribution of Iba-1 positive Cell Area, Cell Perimeter and Area/Perimeter ratio. **A-C**) In the mPFC of heroin experienced rats there was a leftward shift in Cell Area (**A**) and Perimeter (**B**), and a rightward shift of Area/Perimeter ratio (**C**). **D-F**) In the Insula of heroin experienced rats there was a leftward shift in Cell Area (**D**) and Perimeter (**E**), and a rightward shift of Area/Perimeter ratio (**F**). **G-I**) In the Primary Motor cortex of heroin experienced rats there was a leftward shift in Cell Area (**G**) and Perimeter (**H**), but no difference in the Area/Perimeter ratio (**F**). Statistical significance: *p<0.05 and ****p<0.0001 between groups.

In the IC of the heroin experienced group, we observed a leftward shift of both cell area [KS D = 0.18; p<0.0001] and perimeter [KS D = 0.17; p<0.0001] that was maintained until the 87th and 85th percentile of the two cumulative distributions respectively (**Figure 5D, E**). Also in this case, the smaller area and perimeter associated with a larger area/perimeter ratio in the heroin experienced rats [KS D = 0.11; p<0.05] (**Figure 5F**), was due to larger decrease in cell perimeter (−10.1%) than in cell area (−6.5%)

In M1, similar to the other two regions analyzed, we observed left-ward shift of both microglial cells area [KS D = 0.27; p<0.0001, −30% on average] and perimeter [KS D = 0.3; p<0.0001, −33%] (**Figure 5G, H**). However, in this case, the smaller area and perimeter was not associated with changes in the area/perimeter ratio [KS D = 0.07; p>0.05] (**Figure 5I**).

For both Iba-1 and PV positive cells, the morphological parameters’ distributions were similar across groups and are reported in **Supplementary Figures S6 and S7**.

In summary, the population of microglial cells sampled in both groups contained a wide range of sizes and morphologies (**Figure 6**), but while control rats were enriched in highly ramified cell phenotypes (**Figure 6C**), heroin experienced rats contained smaller and less ramified cells (**Figure 6A**).

**FIGURE 6:**
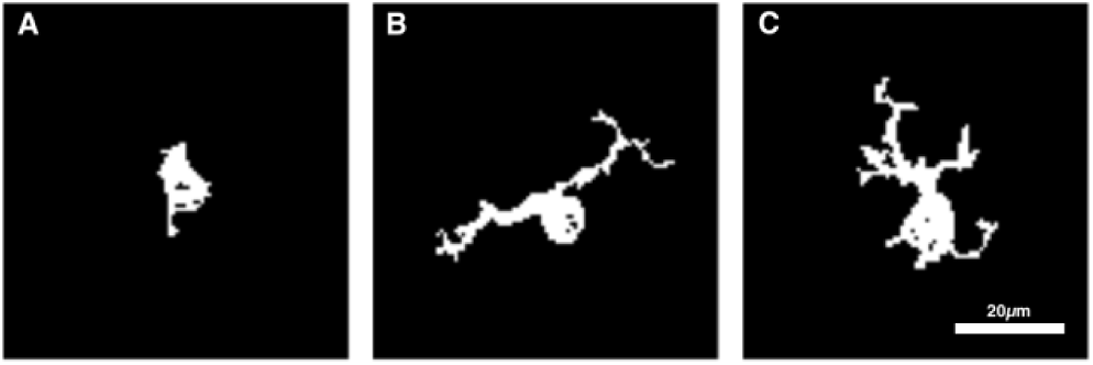
Examples of automatically segmented microglial cells showing low (**A**), medium (**B**) and highly (**C**) ramified phenotypes, scale bar 20 μm.

## 1. DISCUSSION

Our results demonstrate a significant escalation of heroin intake and enhanced motivation for the drug following protracted LgA self-administration. These features reflect some of the primary DSM criteria for OUD diagnosis (i.e., the substance is often taken in larger amounts and over a larger period than intended) (Edwards and Koob, 2013). Escalation and enhanced motivation for heroin following LgA exposure were associated with a reduction of GMV in cortical and subcortical regions shown to being part of the addiction neurocircuitry (Koob and Volkow, 2016). Moreover, consistent with observations in heroin dependent patients (Lin et al., 2018; Qiu et al., 2013; Schmidt et al., 2021; Shi et al., 2020; Wang et al., 2012), we found reduction of GMV extending from the Cg1 dorsally, to the IL ventrally (Paxinos and Watson, 1998). These structures are functionally, and to a large extent anatomically, homologous to the dorsolateral prefrontal cortex and anterior cingulate cortex in primates (Seamans et al., 2008; Uylings et al., 2003). In addition, as previously described in human opioid abusers, we observed GMV reduction in the insular cortex (Bach et al., 2019; Bach et al., 2021). Our results expand these findings with human as our longitudinal approach and the use of an age matched control group support two major conclusions: first, changes in GMV in these regions appears to be the consequence of heroin exposure rather than being a predisposing factor to heroin abuse; second, due to the tightly controlled environmental conditions used with the laboratory animal experiments it is conceivable to argue that heroin exposure alone is adequate to induce these changes in GMV, thereby eliminating the influence of common confounding factors (i.e, patient history, lifestyle etc.) intrinsically associated with clinical studies. Nonetheless, the contribution of environmental factors in affecting GMV changes observed in OUD patients should not be overlooked.

To further explore the link between heroin consumption and brain structural alterations, we also ran correlation analyses between behavioral readouts and GMV changes. We found a negative correlation between the total heroin intake and escalation of drug consumption with GMV reduction in cortical regions and in the striatum. This finding further strengthens the hypothesis that reduced GMV is a consequence of heroin consumption. However, this does not rule out the possibility that diminished GMV could contribute to shaping the progression of OUD. For instance, the mPFC modulates impulse controls and the immaturity or damage of this area has been associated with poor control inhibition and the undertaking of risky behavior, including drug binging (Perry et al., 2011). One could argue that chronic heroin consumption reduces GMV in fronto-cortical regions, which in turn results in a poor behavioral control that further facilitates binging episodes, eventually leading to escalation of heroin intake. In a self-powered vicious cycle, enhanced drug consumption may then contribute to further reduction of GMV. In other words, it is possible that heroin consumption leads to the reduction of GMV which would then promote further heroin seeking. When correlational analysis was applied to heroin extinction/seeking data, a significant link with GMV reduction was found in the insula and DS. It is tempting to speculate that reduced GMV in the insula may reflect changes in interoceptive perception of heroin (in the case of priming) and/or in drug abstinence (Droutman et al., 2015). Also, structural changes in the DS may be linked to aberrant habit learning and promotion of relapse-like behavior (Everitt and Robbins, 2013).

To gain further insights on the morphological changes occurred following heroin we expanded our investigation to determine possible cellular adaptations associated with GMV reduction.

In a preliminary analysis of an ongoing wide transcriptomic investigation in HS rats trained for LgA heroin self-administration, we observed that drug exposure induced significant expression changes in a number of genes associated with immune modulation and microglia activity in the prelimbic cortex (see **Supplementary Table S1**). Notably, the prelimbic cortex is one of the subregions in which structural changes were observed in our MRI study. In addition to this finding, previous work studying the cellular correlate of GMV variations showed multiple potential mechanisms, including changes in the density and size of neuronal and glial cells, but also remodeling of dendritic spines. For instance, after heart failure, a decreased number of neurons and increased microglia were found in the same region where GM concentration was reduced (Bach et al., 2019). However, in other cases, changes in voxel-based morphometry were mostly explained by changes in dendritic spine density or cell clustering rather than in cell death or proliferation (Asan et al., 2021; Keifer et al., 2015).

Inspired by these results we decided to carry out an immunohistochemistry analysis of neurons and microglia in two regions (mPFC and IC) in which the longitudinal MRI experiment showed decreased GMV, and one region (M1) in which GMV was unaltered.

Results showed that heroin self-administration did not affect neuronal and microglia cell density except for an increased number of PV neurons in the motor cortex. In the insula, we observed a rightward shift of both area and perimeter distributions in heroin-exposed animals relative to controls, indicating an increase in the size of these neurons. Conversely, no significant changes were detected in the mPFC and M1. When we looked more in depth at the microglia, we found that heroin self-administration was associated with an overall shrinkage of these cells and an increased area/perimeter ratio in the IC and in the mPFC. This indicated a possible retraction of microglial processes and a shift towards a more amoeboid reactive state of these cells. Interestingly, in M1, where GMV was not altered by heroin, the shrinkage of microglia was not accompanied by a change in the area/perimeter ratio suggesting a different kind of morphological reshaping and perhaps of reactivity. De Santis and colleagues demonstrated that following alcohol drinking, the switch of microglia morphology to a less ramified shape decreased the extracellular space complexity in the grey matter, increasing mean diffusivity and promoting neurotransmitter diffusion (De Santis et al., 2020). Likewise, the increased area/perimeter ratio observed in in our heroin exposed rats may be due to a less ramified morphology of the microglia resulting in a decreased extracellular space complexity and subsequent enhancement of neurotransmitter diffusion from release sites.

Microglia are specialized resident immune cells of the central nervous system, which upon insult can react to mediate pro- or anti-inflammatory processes (Paolicelli et al., 2022). A change in the microglial state is accompanied by a morphological transformation, from a highly ramified phenotype with thin processes extending far from the small soma, to a less ramified shape with shorter and dynamic processes and a larger amoeboid-like appearance (Stence et al., 2001). Even though we did not directly assess the number and length of microglial process ramifications, the increase in the cell area/perimeter ratio is compatible with a reduction in their complexity. Reabsorption of microglial processes and shift from a highly ramified to a less ramified shape is consistent with a possible inflammatory state in heroin treated animals relative to controls in the mPFC and Insula. The presence of a pro-inflammatory state associated with heroin escalation still needs to be demonstrated with specific experiments (Davis et al., 2017; Stence et al., 2001), however, this would be in line with the pharmacological effects of heroin’s metabolites. Heroin is de-acetylated to monoacetyl-morphine and then to morphine, which is converted to morphine-6-glucuronide (M6G) and morphine-3-glucuronide (M3G) (Milella et al., 2023; Rook et al., 2006). While M6G contributes to the rewarding effects of heroin, M3G has no affinity for opioid receptors, but binds and activates the toll-like receptor 4 (TL4) / myeloid differentiation factor 2 (MD2) complex on microglia surface (Green et al., 2022), which in turn leads to the release of pro-inflammatory factors like TNFα and IL-1β. This effect of M3G would be consistent with an inflammatory state associated with the increased area/perimeter ratio that we observed. Noteworthy, in our preliminary RNAseq analysis we found genes such as *Il1r1*, that encodes IL-1β cognate receptor, and *Nfkbib*, a modulator of the TNF signaling pathway (Yazdi & Ghoreschi, 2016; Fields et al., 2019) among the genes which expression was altered by heroin. It is also possible that the stimulation of microglia reactivity by the heroin metabolite M3G contributed to the escalation of heroin intake. Indeed, it was reported that blockade of glial reactivity reduced morphine induced release of dopamine in the ventral striatum (Bland et al., 2009), possibly through a TLR4 dependent mechanism, as TLR4 knock out mice do not develop opioid place preference (Hutchinson et al., 2012). Consistently, the blockade of TLR4 by (+)-naltrexone decreased operant self-administration of remifentanil (Yue et al., 2020), though the selectivity of this effect has been argued (Tanda et al., 2016). In addition to opioid reward, activation of TLR4 plays a role in the development of opioid tolerance (Eidson and Murphy, 2013). It is thus possible that by decreasing reward and increasing tolerance, the reactive state of microglia contributed to the escalation of heroin intake.

The under-representation of female subjects has been a bias in both clinical and preclinical research since many years. To avoid this bias, in this study we adopted a sex matched approach. However, while a sex-matched experimental design allowed us to generalize between male and female, the sample size of our experiments is insufficient to evaluate if sex differences in response to heroin occurred (Paolicelli et al., 2022).

In conclusion, our data demonstrated that heroin consumption under long access contingency led to region-specific reduction in GMV that correlated with the quantity and escalation of heroin intake. Such reduction was confined to cortical regions known to be consequential in substance use disorder. GMV reduction occurred in the same areas where alterations in microglia morphology were also observed, which suggests that this cell population may contribute to gray matter alterations following heroin.

## Acknowledgement

This work was supported by U01-DA045300 to GH, LSW, DC, PK and RC. The Authors wish to thank Agostino Marchi, Rina Righi and Matteo Valzano for animal care and technical support. AB acknowledges support by the Fondazione Cassa di Risparmio di Torino (CRT), R.F. 2019.0610.

## Author Contributions

NC and RC ideated the project, NC coordinated the implementation of this work, ran behavioral experiment, and analyzed data, RC supervised the project. ST acquired scans and analyzed MRI data. VL ran self-administration sessions, and immunofluorescence. GS acquired MRI scans, LdV supervised immunofluorescence, SA, AK, JM, and GH provided preliminary RNAseq data, LCSW supervised HS rats breeding, LS and MU sampled tissues, AB supervised MRI acquisitions and analyses. NC, ST, LDV and RC wrote the manuscript. All authors contributed to the article and approved the submitted version.

## REFERENCES

Ahmed, S.H., Koob, G.F., 1998. Transition from moderate to excessive drug intake: change in hedonic set point. Science 282, 298–300.

Allen, C., Kuhn, B.N., Cannella, N., Crow, A.D., Roberts, A.T., Lunerti, V., Ubaldi, M., Hardiman, G., Solberg Woods, L.C., Ciccocioppo, R., Kalivas, P.W., Chung, D., 2021. Network-Based Discovery of Opioid Use Vulnerability in Rats Using the Bayesian Stochastic Block Model. Front Psychiatry 12, 745468.

Asan, L., Falfan-Melgoza, C., Beretta, C.A., Sack, M., Zheng, L., Weber-Fahr, W., Kuner, T., Knabbe, J., 2021. Cellular correlates of gray matter volume changes in magnetic resonance morphometry identified by two-photon microscopy. Sci Rep 11, 4234.

Ashburner, J., Friston, K.J., 2000. Voxel-based morphometry--the methods. Neuroimage 11, 805–821.

Bach, P., Frischknecht, U., Klinkowski, S., Bungert, M., Karl, D., Vollmert, C., Vollstadt-Klein, S., Lis, S., Kiefer, F., Hermann, D., 2019. Higher Social Rejection Sensitivity in Opioid-Dependent Patients Is Related to Smaller Insula Gray Matter Volume: A Voxel-Based Morphometric Study. Soc Cogn Affect Neurosci 14, 1187–1195.

Bach, P., Frischknecht, U., Reinhard, I., Bekier, N., Demirakca, T., Ende, G., Vollstadt-Klein, S., Kiefer, F., Hermann, D., 2021. Impaired working memory performance in opioid-dependent patients is related to reduced insula gray matter volume: a voxel-based morphometric study. Eur Arch Psychiatry Clin Neurosci 271, 813–822.

Bellesi, M., de Vivo, L., Chini, M., Gilli, F., Tononi, G., Cirelli, C., 2017. Sleep Loss Promotes Astrocytic Phagocytosis and Microglial Activation in Mouse Cerebral Cortex. J Neurosci 37, 5263–5273.

Bland, S.T., Hutchinson, M.R., Maier, S.F., Watkins, L.R., Johnson, K.W., 2009. The glial activation inhibitor AV411 reduces morphine-induced nucleus accumbens dopamine release. Brain Behav Immun 23, 492–497.

Davis, B.M., Salinas-Navarro, M., Cordeiro, M.F., Moons, L., De Groef, L., 2017. Characterizing microglia activation: a spatial statistics approach to maximize information extraction. Sci Rep 7, 1576.

de Guglielmo, G., Kallupi, M., Sedighim, S., Newman, A.H., George, O., 2019. Dopamine D(3) Receptor Antagonism Reverses the Escalation of Oxycodone Self-administration and Decreases Withdrawal-Induced Hyperalgesia and Irritability-Like Behavior in Oxycodone-Dependent Heterogeneous Stock Rats. Front Behav Neurosci 13, 292.

de Guglielmo, G., Melis, M., De Luca, M.A., Kallupi, M., Li, H.W., Niswender, K., Giordano, A., Senzacqua, M., Somaini, L., Cippitelli, A., Gaitanaris, G., Demopulos, G., Damadzic, R., Tapocik, J., Heilig, M., Ciccocioppo, R., 2015. PPARgamma activation attenuates opioid consumption and modulates mesolimbic dopamine transmission. Neuropsychopharmacology 40, 927–937.

De Santis, S., Cosa-Linan, A., Garcia-Hernandez, R., Dmytrenko, L., Vargova, L., Vorisek, I., Stopponi, S., Bach, P., Kirsch, P., Kiefer, F., Ciccocioppo, R., Sykova, E., Moratal, D., Sommer, W.H., Canals, S., 2020. Chronic alcohol consumption alters extracellular space geometry and transmitter diffusion in the brain. Sci Adv 6, eaba0154.

Droutman, V., Read, S.J., Bechara, A., 2015. Revisiting the role of the insula in addiction. Trends Cogn Sci 19, 414–420.

Edwards, S., Koob, G.F., 2013. Escalation of drug self-administration as a hallmark of persistent addiction liability. Behav Pharmacol 24, 356–362.

Eidson, L.N., Murphy, A.Z., 2013. Blockade of Toll-like receptor 4 attenuates morphine tolerance and facilitates the pain relieving properties of morphine. J Neurosci 33, 15952–15963.

Everitt, B.J., Robbins, T.W., 2013. From the ventral to the dorsal striatum: devolving views of their roles in drug addiction. Neurosci Biobehav Rev 37, 1946–1954.

Fields, J.K., Günther, S., Sundberg, E.J.. 2019. Structural Basis of IL-1 Family Cytokine Signaling. Front Immunol, 20:10:1412.

Frischknecht, U., Beckmann, B., Heinrich, M., Kniest, A., Nakovics, H., Kiefer, F., Mann, K., Hermann, D., 2011. The vicious circle of perceived stigmatization, depressiveness, anxiety, and low quality of life in substituted heroin addicts. Eur Addict Res 17, 241–249.

Green, J.M., Sundman, M.H., Chou, Y.H., 2022. Opioid-induced microglia reactivity modulates opioid reward, analgesia, and behavior. Neurosci Biobehav Rev 135, 104544.

Hahm, S., Lotze, M., Domin, M., Schmidt, S., 2019. The association of health-related quality of life and cerebral gray matter volume in the context of aging: A voxel-based morphometry study with a general population sample. Neuroimage 191, 470–480.

Hutchinson, M.R., Northcutt, A.L., Hiranita, T., Wang, X., Lewis, S.S., Thomas, J., van Steeg, K., Kopajtic, T.A., Loram, L.C., Sfregola, C., Galer, E., Miles, N.E., Bland, S.T., Amat, J., Rozeske, R.R., Maslanik, T., Chapman, T.R., Strand, K.A., Fleshner, M., Bachtell, R.K., Somogyi, A.A., Yin, H., Katz, J.L., Rice, K.C., Maier, S.F., Watkins, L.R., 2012. Opioid activation of toll-like receptor 4 contributes to drug reinforcement. J Neurosci 32, 11187–11200.

Kallupi, M., Carrette, L.L.G., Kononoff, J., Solberg Woods, L.C., Palmer, A.A., Schweitzer, P., George, O., de Guglielmo, G., 2020. Nociceptin attenuates the escalation of oxycodone self-administration by normalizing CeA-GABA transmission in highly addicted rats. Proc Natl Acad Sci U S A 117, 2140–2148.

Keifer, O.P., Jr., Hurt, R.C., Gutman, D.A., Keilholz, S.D., Gourley, S.L., Ressler, K.J., 2015. Voxel-based morphometry predicts shifts in dendritic spine density and morphology with auditory fear conditioning. Nat Commun 6, 7582.

Koob, G.F., Volkow, N.D., 2016. Neurobiology of addiction: a neurocircuitry analysis. Lancet Psychiatry 3, 760–773.

Kuhn, B.N., Cannella, N., Crow, A.D., Roberts, A.T., Lunerti, V., Allen, C., Nall, R.W., Hardiman, G., Woods, L.C.S., Chung, D., Ciccocioppo, R., Kalivas, P.W., 2022. Novelty-induced locomotor behavior predicts heroin addiction vulnerability in male, but not female, rats. Psychopharmacology (Berl) 239, 3605–3620.

Lin, H.C., Wang, P.W., Wu, H.C., Ko, C.H., Yang, Y.H., Yen, C.F., 2018. Altered gray matter volume and disrupted functional connectivity of dorsolateral prefrontal cortex in men with heroin dependence. Psychiatry Clin Neurosci 72, 435–444.

Liu, H., Hao, Y., Kaneko, Y., Ouyang, X., Zhang, Y., Xu, L., Xue, Z., Liu, Z., 2009. Frontal and cingulate gray matter volume reduction in heroin dependence: optimized voxel-based morphometry. Psychiatry Clin Neurosci 63, 563–568.

Milella, M.S., D’Ottavio, G., De Pirro, S., Barra, M., Caprioli, D., Badiani, A., 2023. Heroin and its metabolites: relevance to heroin use disorder. Transl Psychiatry 13, 120.

NIDA, 2023. Drug Overdose Death Rates.

Paolicelli, R.C. et al., 2022. Microglia states and nomenclature: A field at its crossroads. Neuron. 2022 Nov 2;110(21):3458–3483.

Paxinos, G., Watson, C., 1998. The rat brain.

Perry, J.L., Joseph, J.E., Jiang, Y., Zimmerman, R.S., Kelly, T.H., Darna, M., Huettl, P., Dwoskin, L.P., Bardo, M.T., 2011. Prefrontal cortex and drug abuse vulnerability: translation to prevention and treatment interventions. Brain Res Rev 65, 124–149.

Puigdollers, E., Domingo-Salvany, A., Brugal, M.T., Torrens, M., Alvaros, J., Castillo, C., Magri, N., Martin, S., Vazquez, J.M., 2004. Characteristics of heroin addicts entering methadone maintenance treatment: quality of life and gender. Subst Use Misuse 39, 1353–1368.

Qiu, Y.W., Jiang, G.H., Su, H.H., Lv, X.F., Tian, J.Z., Li, L.M., Zhuo, F.Z., 2013. The impulsivity behavior is correlated with prefrontal cortex gray matter volume reduction in heroin-dependent individuals. Neurosci Lett 538, 43–48.

Richardson, N.R., Roberts, D.C., 1996. Progressive ratio schedules in drug self-administration studies in rats: a method to evaluate reinforcing efficacy. J Neurosci Methods 66, 1–11.

Rook, E.J., Huitema, A.D., van den Brink, W., van Ree, J.M., Beijnen, J.H., 2006. Pharmacokinetics and pharmacokinetic variability of heroin and its metabolites: review of the literature. Curr Clin Pharmacol 1, 109–118.

Schmidt, A., Vogel, M., Baumgartner, S., Wiesbeck, G.A., Lang, U., Borgwardt, S., Walter, M., 2021. Brain volume changes after long-term injectable opioid treatment: A longitudinal voxel-based morphometry study. Addict Biol 26, e12970.

Schnabel, J.A., Tanner, C., Castellano-Smith, A.D., Degenhard, A., Leach, M.O., Hose, D.R., Hill, D.L., Hawkes, D.J., 2003. Validation of nonrigid image registration using finite-element methods: application to breast MR images. IEEE Trans Med Imaging 22, 238–247.

Seamans, J.K., Lapish, C.C., Durstewitz, D., 2008. Comparing the prefrontal cortex of rats and primates: insights from electrophysiology. Neurotox Res 14, 249–262.

Shaham, Y., Shalev, U., Lu, L., de Wit, H., Stewart, J., 2003. The reinstatement model of drug relapse: history, methodology and major findings. Psychopharmacology (Berl) 168, 3–20.

Shi, H., Liang, Z., Chen, J., Li, W., Zhu, J., Li, Y., Ye, J., Zhang, J., Xue, J., Liu, W., Wang, F., Wang, W., Li, Q., He, X., 2020. Gray matter alteration in heroin-dependent men: An atlas-based magnetic resonance imaging study. Psychiatry Res Neuroimaging 304, 111150.

Smith, S.M., Jenkinson, M., Woolrich, M.W., Beckmann, C.F., Behrens, T.E., Johansen-Berg, H., Bannister, P.R., De Luca, M., Drobnjak, I., Flitney, D.E., Niazy, R.K., Saunders, J., Vickers, J., Zhang, Y., De Stefano, N., Brady, J.M., Matthews, P.M., 2004. Advances in functional and structural MR image analysis and implementation as FSL. Neuroimage 23 Suppl 1, S208–219.

Solberg Woods, L.C., Palmer, A.A., 2019. Using Heterogeneous Stocks for Fine-Mapping Genetically Complex Traits. Methods Mol Biol 2018, 233–247.

Stence, N., Waite, M., Dailey, M.E., 2001. Dynamics of microglial activation: a confocal time-lapse analysis in hippocampal slices. Glia 33, 256–266.

Sun, Y., Liu, L., Feng, J., Yue, W., Lu, L., Fan, Y., Shi, J., 2017. MAOA rs1137070 and heroin addiction interactively alter gray matter volume of the salience network. Sci Rep 7, 45321.

Sun, Y., Zhao, L.Y., Wang, G.B., Yue, W.H., He, Y., Shu, N., Lin, Q.X., Wang, F., Li, J.L., Chen, N., Wang, H.M., Kosten, T.R., Feng, J.J., Wang, J., Tang, Y.D., Liu, S.X., Deng, G.F., Diao, G.H., Tan, Y.L., Han, H.B., Lin, L., Shi, J., 2016. ZNF804A variants confer risk for heroin addiction and affect decision making and gray matter volume in heroin abusers. Addict Biol 21, 657–666.

Tambalo, S., Peruzzotti-Jametti, L., Rigolio, R., Fiorini, S., Bontempi, P., Mallucci, G., Balzarotti, B., Marmiroli, P., Sbarbati, A., Cavaletti, G., Pluchino, S., Marzola, P., 2015. Functional Magnetic Resonance Imaging of Rats with Experimental Autoimmune Encephalomyelitis Reveals Brain Cortex Remodeling. J Neurosci 35, 10088–10100.

Tanda, G., Mereu, M., Hiranita, T., Quarterman, J.C., Coggiano, M., Katz, J.L., 2016. Lack of Specific Involvement of (+)-Naloxone and (+)-Naltrexone on the Reinforcing and Neurochemical Effects of Cocaine and Opioids. Neuropsychopharmacology 41, 2772–2781.

Uylings, H.B., Groenewegen, H.J., Kolb, B., 2003. Do rats have a prefrontal cortex? Behav Brain Res 146, 3–17.

Vendruscolo, L.F., Schlosburg, J.E., Misra, K.K., Chen, S.A., Greenwell, T.N., Koob, G.F., 2011. Escalation patterns of varying periods of heroin access. Pharmacol Biochem Behav 98, 570–574.

Wang, L., Zou, F., Zhai, T., Lei, Y., Tan, S., Jin, X., Ye, E., Shao, Y., Yang, Y., Yang, Z., 2016. Abnormal gray matter volume and resting-state functional connectivity in former heroin-dependent individuals abstinent for multiple years. Addict Biol 21, 646–656.

Wang, X., Li, B., Zhou, X., Liao, Y., Tang, J., Liu, T., Hu, D., Hao, W., 2012. Changes in brain gray matter in abstinent heroin addicts. Drug Alcohol Depend 126, 304–308.

Yazdi, A.S., Ghoreschi, K., 2016. The Interleukin-1 Family, Adv Exp Med Biol 941, 21–29.

Yen, C.N., Wang, C.S., Wang, T.Y., Chen, H.F., Chang, H.C., 2011. Quality of life and its correlates among heroin users in Taiwan. Kaohsiung J Med Sci 27, 177–183.

Yue, K., Tanda, G., Katz, J.L., Zanettini, C., 2020. A further assessment of a role for Toll-like receptor 4 in the reinforcing and reinstating effects of opioids. Behav Pharmacol 31, 186–195.

