## supplementary information for "Long-access heroin self-administration induces region specific reduction of grey matter volume and microglia reactivity in the rat"

### MATERIALS AND METHODS

#### Heroin self-administration training

Surgery

Rats in the heroin experienced group were surgically implanted with an indwelling catheter in the right jugular vein under isoflurane anesthesia (1.5-2.5%). Incisions were made to expose the right jugular vein and the back between the shoulders. A catheter composed of a Micro-Renathane® tube (I.D.=0.020in, O.D.=0.037in; Braintree Scientific) and an MRI-compatible back-mount (Plastic-One) was subcutaneously positioned between these two points. After insertion into the vein, the proximal end of the catheter was anchored to the muscles underlying the vein with surgical silk sutures. The back-mount at the distal end passed under the skin and protruded from the mid-scapular region.

Rats were allowed to recover 1 week before the beginning of the self-administration training. Catheters were flushed with 100 µL/rat of heparinized saline (nadroparin calcium 100 UI/mL) containing 1.0 mg/mL of enrofloxacin after every SA session. In control heroin naive rats, the right jugular vein was exposed and occluded with surgical silk.

Self-administration apparatus

Experiments were conducted in self-administration stations consisting of operant self-administration chambers (Med Associate Inc. St. Alban, USA) enclosed in sound-attenuating, ventilated environmental cubicles. Each chamber was equipped with two retractable levers located on the front panel of the chamber five cm above the grid floor, and a cue light above each lever. A house-light and a tone generator were located on the opposite wall. To deliver heroin, one end of a tygon tube, enclosed into a metallic tether, was connected to the catheter whereas the other extreme was connected to an infusion pump via a swivel. Activation of the pump resulted in a delivery of 0.1 ml of fluid. An IBM compatible computer controlled the delivery of heroin solution and the recording of data.

Food-reinforced pre-training of operant responding

Food pre-training was conducted in a separate set of self-administration boxes equipped with a receptacle to deliver food pellets between the levers. Rats were initially trained to operant self-administration in four 30-minute food self-administration sessions. Sessions started with the insertion of the right lever. Lever pressing under fixed ratio 1 (FR1) schedule of reinforcement activated the pellet dispenser and delivered one food pellet (TestDiet: catalog #181156) in a receptacle located on the left side of the lever. Pellet delivery was associated with the contingent illumination of the house light that remained on for 5 seconds. During the 4 days of food training, food availability in the home cage was limited to 5g/day of standard chow pellets.

**RNAseq Analysis**

Brain dissection, RNA isolation and library preparation

The prelimbic cortex was micro-dissected at -20 °C using Leica cryostat (CM1900) using the Paxinos and Watson rat brain atlas 6$^{th}$ edition and following the stereotaxic coordinates of Bregma 3.24mm and Interaural 12.24mm. Punches were taken from the exposed coronal section using 19-gauge blunt needles. Samples were transferred to clean RNase/DNase free Eppendorf tubes and stored at -80 °C for further processing.

Total RNA was isolated from the prelimbic cortex of 16 HS rat brain randomly collected from the tissue bank generated in our earlier Allen et al. (2021) study [19] (4 heroin naïve male rats, 6 males and 6 female rats subjected to LgA heroin self-administration) using Qiagen miRNeasy kit following the manufacturer's guide.

RNA samples were checked for integrity using Agilent TapeStation system and only RNA samples of RNA integrity number (RIN) $\geq$ 7 were used for library preparation. Library preparation was performed using a starting volume of 100 ng RNA and the Illumina® Stranded Total RNA Prep, Ligation with Ribo-Zero Plus kit following the manufacturer protocol. Libraries were checked for quality using the Agilent TapeStation system, index balance on MiSeq Nano, sequenced using NovaSeq 6000 platform. Paired end sequencing was performed to a read length of 2X 125bp and a total of 120M reads per sample.

Bioinformatics and data analysis

The quality of sequencing was assessed using Fastqc [1] and the results were visualised by multiqc. Adapters sequences were removed using cutadapt before alignment to the rat genome (mRatBN7.2) using STAR [2]. Counts were generated HTSeq [3], and differential expression analysis was performed using DESeq2 [3]. Sex was incorporated as covariate for the statistical analysis performed by DESeq2, which generated data output including normalised expression values, log fold change (log FC), p value, and adjusted p value. Differential gene expression was determined using FDR $\leq$ 0.4 and log FC higher than or lower than $\pm$ 0.3. Our methodology follows the premise that transcripts are sorted according to their q-value, which is the smallest false discovery rate (FDR) at which the transcript is called significant. FDR is the expected fraction of false positive tests among significant tests. We have shown in recent studies of human breast and prostate cancer [17,18] that an FDR of 0.4 as set by the DEseq2 program permits a sensitive analysis at a systems level of genes that are relevant to the underlying biology being studied.

**Supplementary Table S1**

| **Gene expression in the prelimbic cortex of heroin experienced rats compared to naive** | | |
|---|---|---|
| Gene Symbol | Log Fold Change | Description |
| *Mif* | 0.49 | Mif/Cd74 is a proinflammatory signalling axis found to be involved in highly activated microglia in the ageing rodent brain. Cd74 have been reported to be highly expressed in M1 microglia in the hippocampus following neuronal damage and in glioblastoma. Mif binds its cognate receptor (Cd74) to regulate innate and adaptive immunity [4]. |
| *Cd74* | 1.59 | |
| *Il1r1* | -0.85 | Is a cytokine receptor for IL-1. It is an important mediator of immune signalling induced by cytokines, namely NFκB signalling pathway [5]. |
| *Il6r* | -0.63 | Also known as Il6ra plays an important role in neuroimmune signalling and is selectively expressed in microglia [6]. |
| *Nr4a1* | -0.51 | Is a pleiotropic regulator, in the innate and adaptive immune signalling, that controls microglial activation by preventing overactivation [7]. |
| *Fas* | 0.37 | Fas, also known as Cd95 is a member of the TNF receptor superfamily that can trigger varieties of cellular responses including apoptosis and inflammatory response [8]. |
| *Nfkbib* | 0.48 | Found to be part of the anti-inflammatory transcripts differentially regulated by active microglia isolated from the brain of BALB/cJ$^{Fms\text{-}EGFP/-}$ mice [9]. |
| *Ccr5* | -0.37 | Ccr5 is a gene identified to be specific for microglia [9]. |
| *Sod3* | 0.41 | Play a key role in the NFκB immune signalling pathway as a regulator of transcription [10]. |
| *Nfkb1* | -0.36 | Is a subunit of constituting the transcription factor NFκB which is known for its role in modulating immune responses [11]. |
| *Jund* | -0.37 | Is a transcription factor in the AP-1 family that is known to regulate immune responses via IL-1β [12]. |
| *Cebpb* | -1.05 | Is a transcription factor that regulates various cellular functions besides immune signalling. Cebpb targets numerous pro-inflammatory genes and is known to promote inflammation [13]. |
| *Casp12* | -0.61 | Plays an important role in the regulation of inflammation as a negative regulator of pro-inflammatory cytokines [14]. |
| *Usp18* | 1.32 | Plays a critical role in the immune signalling, especially NFκB signalling pathway, and has been found to regulate microglia state of activity [15]. |
| *Otud1* | -0.42 | Is involved in the regulation of several immune responses including TNF-α/ NFκB [16]. |

### Supplementary Figures

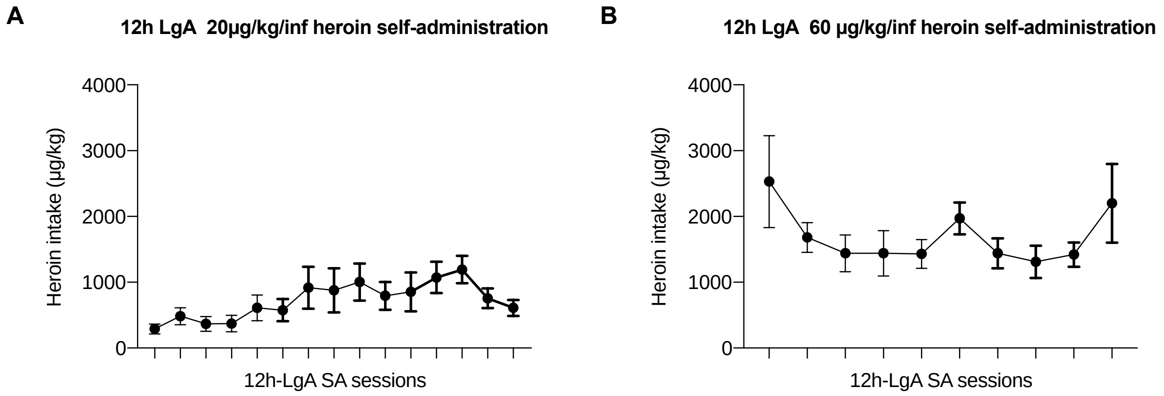


**Figure S1:** Heroin self-administration data of rats used for the histochemical analyses. Self-administration methods were identical to that described in the main text, except that these rats instead of receiving food pre-training were initially acquainted to LgA 20 µg/kg/inf followed by 60 µg/kg/inf that correspond to the dose used in the MRI experiment. **A**) Total daily intake of 20µg/kg/infusion of heroin self-administered during 12h LgA sessions. ANOVA found an overall effect of time [$F(14,70) = 2.54$; $p < 0.01$] indicating acquisition of heroin self-administration. **B)** when rats were switched to 60 µg/kg/infusion the total daily intake increased and remained constant over time $F(9,45) = 1.63$; $p > 0.05$]. Data are presented as mean ± SEM.

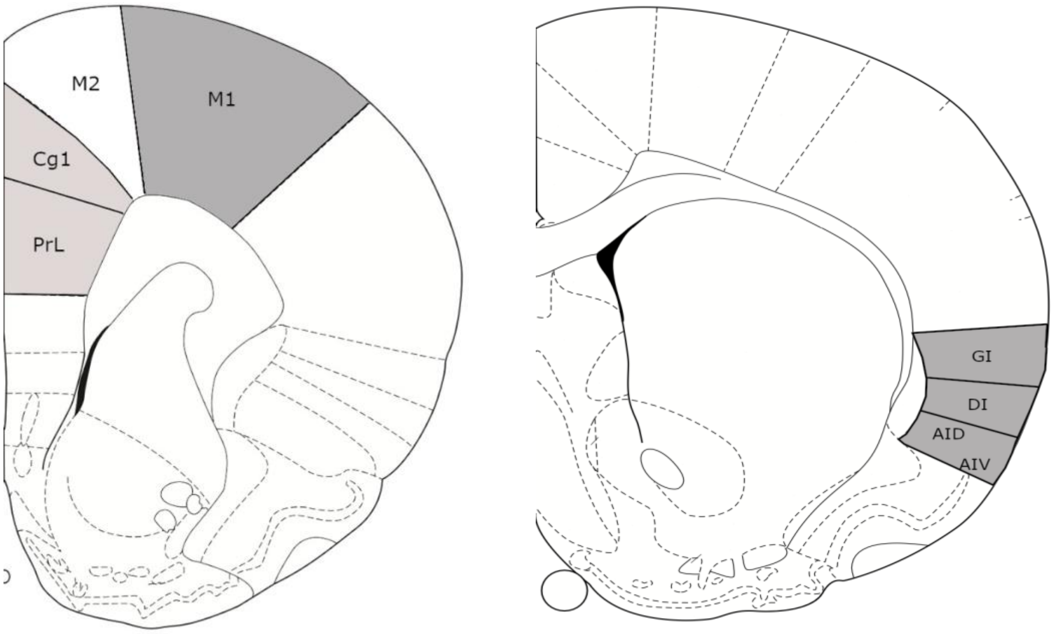

**Figure S2:** schematic representation of brain cortical areas that were considered as mPFC (Cg1 + PrL), primary motor cortex (M1), and Insula (GI + DI + AID/AIV). Pictures were adapted from Paxinos & Watson Rat Brain Atlas (1998).

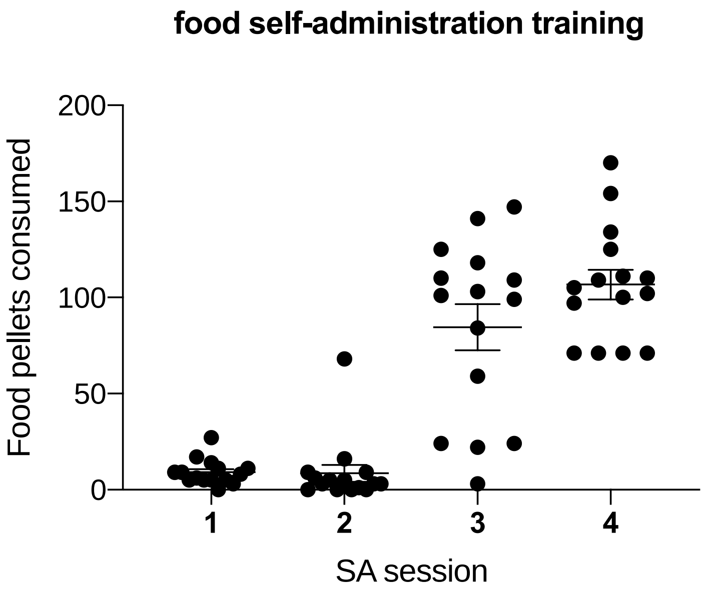


**Figure S3:** Rats were initially trained to operant lever responding during four days of food pretraining. The number of reinforced responses passed from 9.0 ± 1.7 (min = 0, max = 27) on the first day to 106.7 ± 7.7 (min = 71, max = 170) on the fourth day, indicating the acquisition of SA behavior as confirmed by ANOVA [F(3, 14)=66.7; p<0.0001].

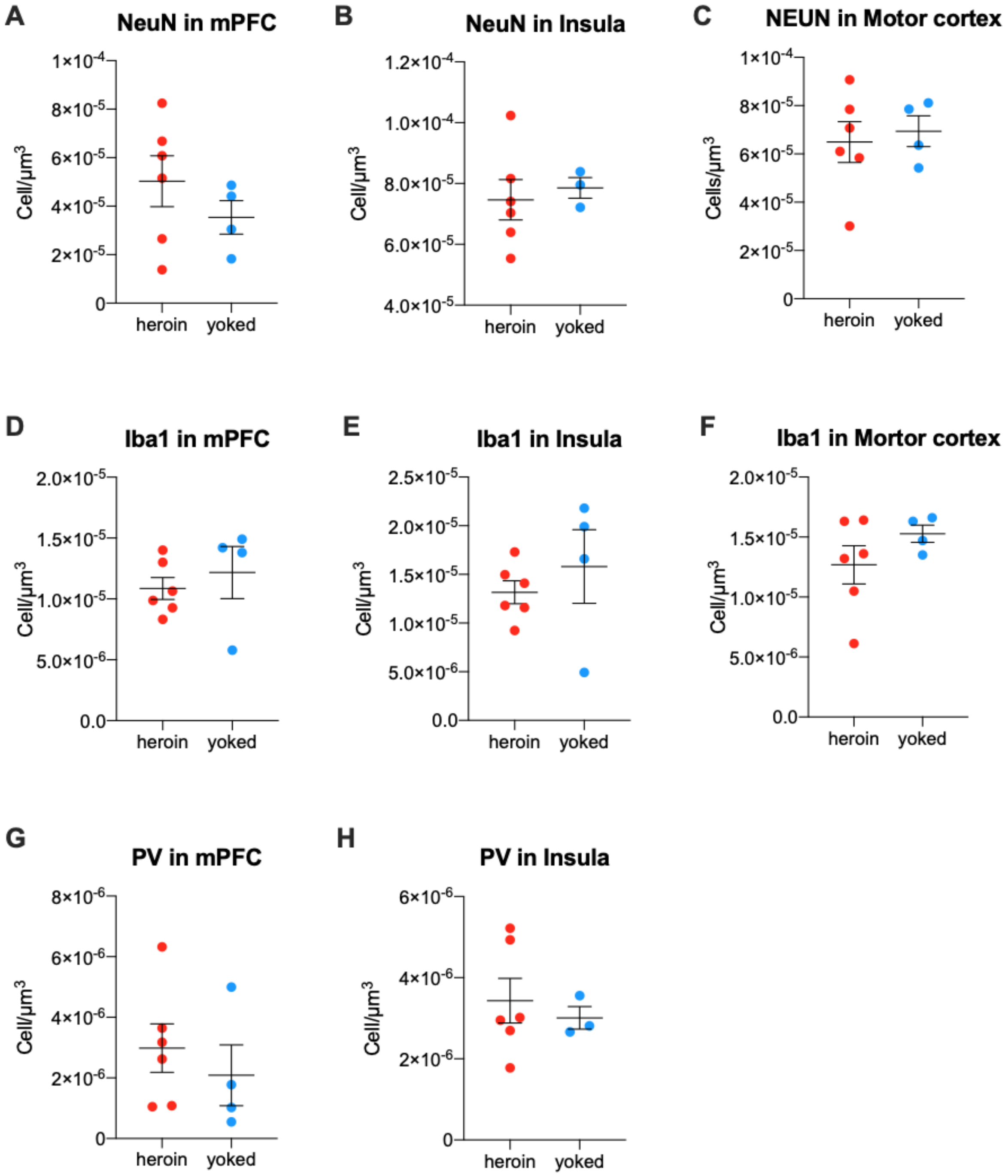


**Figure S4:** Comparison of Neun (**A-C**), PV (**D-F**), and Iba-1 (**G, H**) Cell Density between heroin experienced and heroin yoked rats in mPFC (left column of graphs), Insula (central column) and Motor cortex (right column). Graph **H** reports only three yoked data points because images from one rat could not be acquired for technical issues. data yoked. Whiskers represent mean ± SEM.

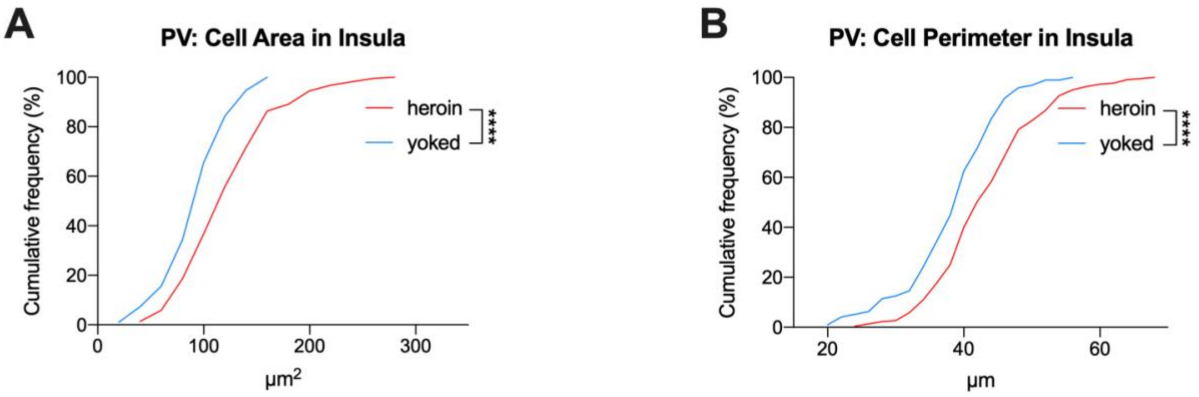


**Figure S5**: In the insula of heroin-experienced rats there was a rightward shift of cumulative PV positive cell Area and Perimeter distributions. Statistical significance: ****p<0.0001.

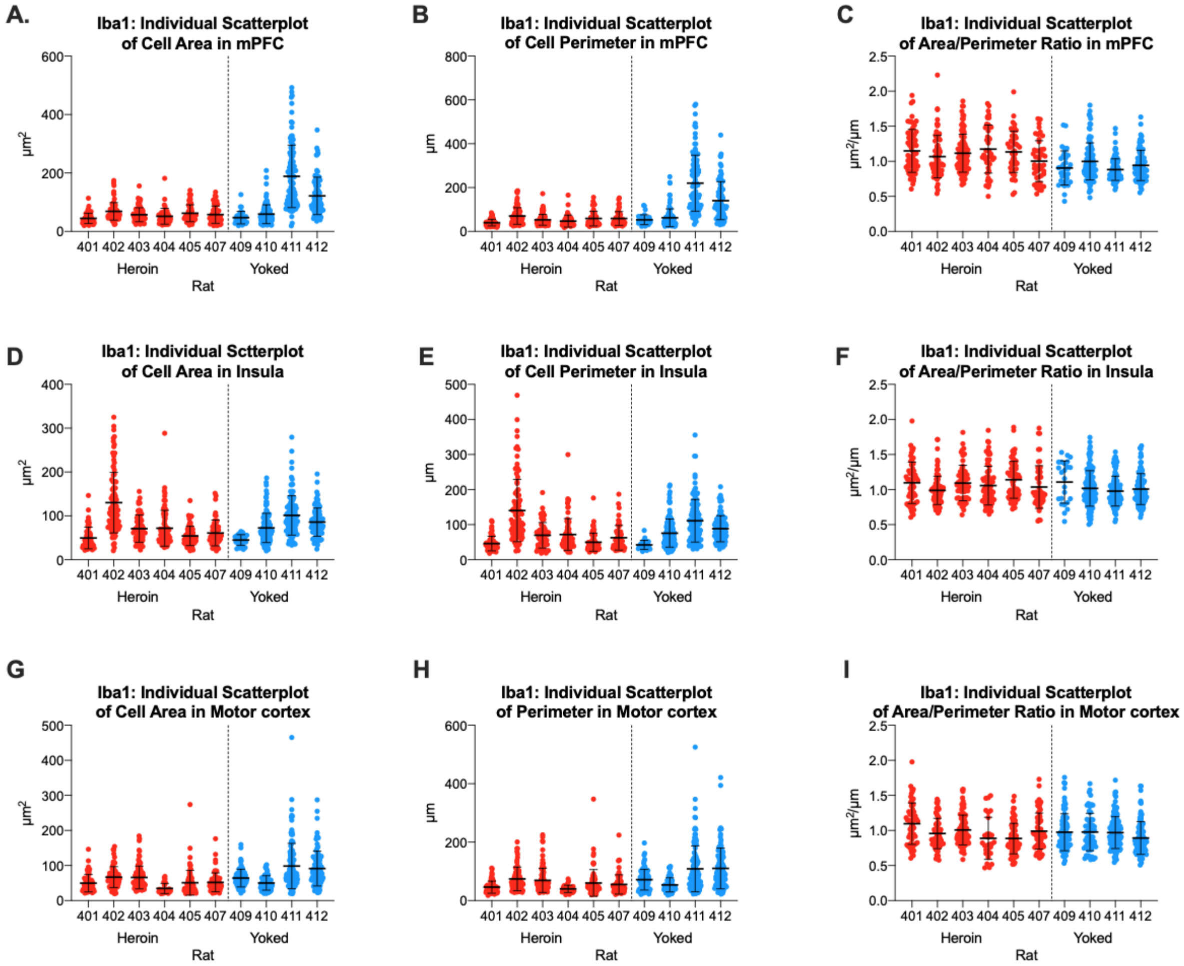


**Figure S6:** Individual scatter plot of Cell Area (left column graphs), Cell Perimeter (central column) and Area/Perimeter ratio (right column) of Iba-1 positive cells in mPFC (**A-C**), Insula (**D-F**) and Motor cortex (**G-I**). Whiskers represent mean ± SEM.

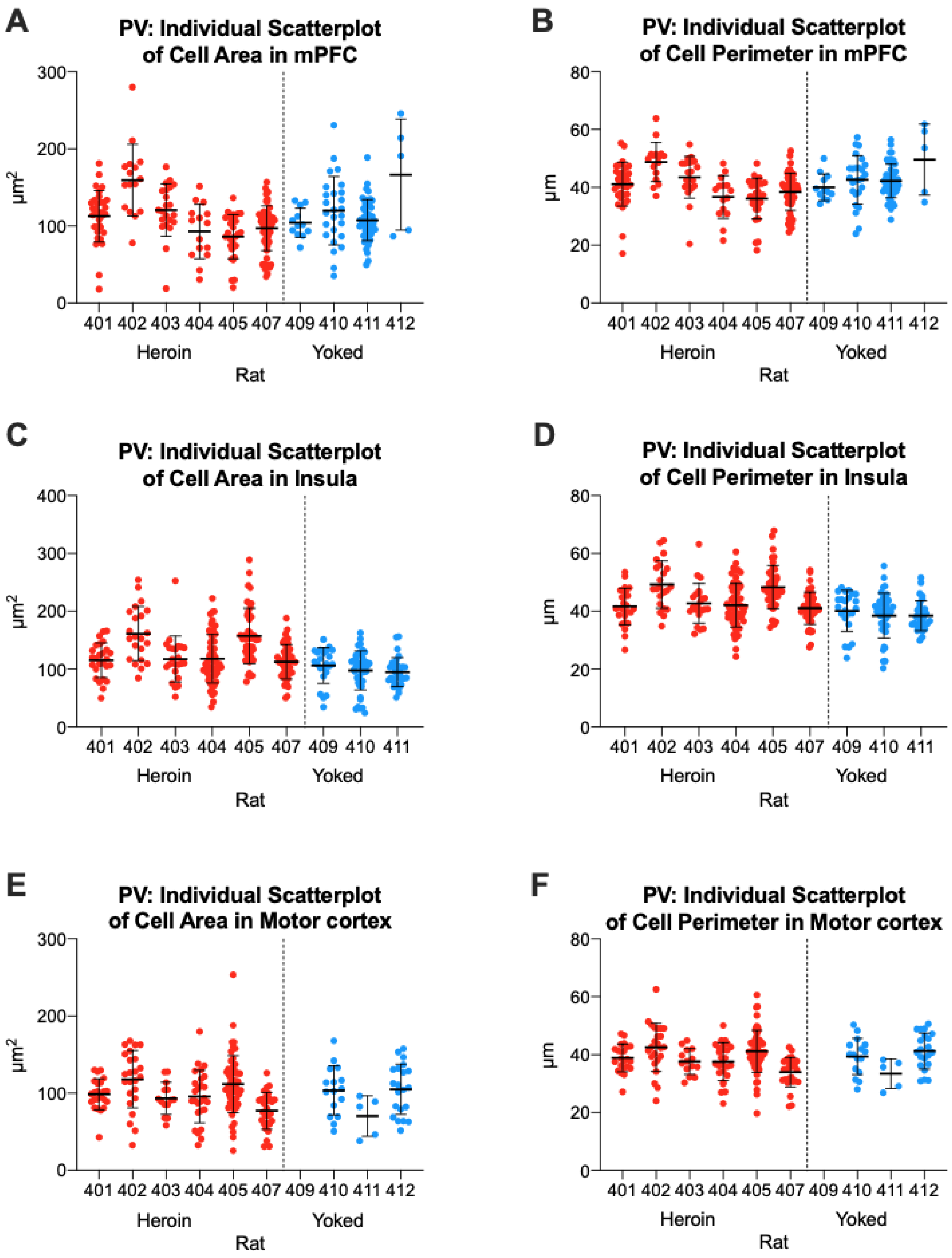


**Figure S7:** Individual scatter plot of Cell Area (left column graphs) and Cell Perimeter (right column) of PV positive cells in mPFC (**A, B**), Insula (**C, D**) and Motor cortex (**E, F**). Whiskers represent mean ± SEM.

**Supplementary References**


1. Lo, C.-C. and P.S. Chain, *Rapid evaluation and quality control of next generation sequencing data with FaQCs.* BMC bioinformatics, 2014. **15**(1): p. 1-8.
2. Dobin, A., et al., *STAR: ultrafast universal RNA-seq aligner.* Bioinformatics, 2013. **29**(1): p. 15-21.
3. Wen, G. *A simple process of RNA-sequence analyses by Hisat2, Htseq and DESeq2*. in *Proceedings of the 2017 international conference on biomedical engineering and bioinformatics*. 2017.
4. Jin, C., et al., *A unique type of highly-activated microglia evoking brain inflammation via Mif/Cd74 signaling axis in aged mice.* Aging and disease, 2021. **12**(8): p. 2125.
5. Liu, X., et al., *Cell-type-specific interleukin 1 receptor 1 signaling in the brain regulates distinct neuroimmune activities.* Immunity, 2019. **50**(2): p. 317-333. e6.
6. Hammond, T.R., et al., *Single-cell RNA sequencing of microglia throughout the mouse lifespan and in the injured brain reveals complex cell-state changes.* Immunity, 2019. **50**(1): p. 253-271. e6.
7. Rothe, T., et al., *The nuclear receptor Nr4a1 acts as a microglia rheostat and serves as a therapeutic target in autoimmune-driven central nervous system inflammation.* The Journal of Immunology, 2017. **198**(10): p. 3878-3885.
8. Choi, C. and E.N. Benveniste, *Fas ligand/Fas system in the brain: regulator of immune and apoptotic responses.* Brain Research Reviews, 2004. **44**(1): p. 65-81.
9. Vincenti, J.E., et al., *Defining the microglia response during the time course of chronic neurodegeneration.* Journal of virology, 2016. **90**(6): p. 3003-3017.
10. Laurila, J.P., et al., *SOD3 reduces inflammatory cell migration by regulating adhesion molecule and cytokine expression.* PloS one, 2009. **4**(6): p. e5786.
11. Nennig, S. and J. Schank, *The role of NFkB in drug addiction: beyond inflammation.* Alcohol and Alcoholism, 2017. **52**(2): p. 172-179.
12. Diaz-Cañestro, C., et al., *AP-1 (activated protein-1) transcription factor JunD regulates ischemia/reperfusion brain damage via IL-1β (interleukin-1β).* Stroke, 2019. **50**(2): p. 469-477.
13. Kapadia, R., et al., *Decreased brain damage and curtailed inflammation in transcription factor CCAAT/enhancer binding protein β knockout mice following transient focal cerebral ischemia.* Journal of neurochemistry, 2006. **98**(6): p. 1718-1731.
14. Garcia de la Cadena, S. and L. Massieu, *Caspases and their role in inflammation and ischemic neuronal death. Focus on caspase-12.* Apoptosis, 2016. **21**: p. 763-777.
15. Xiang, J., et al., *USP18 overexpression protects against focal cerebral ischemia injury in mice by suppressing microglial activation.* Neuroscience, 2019. **419**: p. 121-128.
16. Lin, J., et al., *Herpesvirus latent infection promotes stroke via activating the OTUD1/NF-κB signaling pathway.* Aging (Albany NY), 2023. **15**(17): p. 8976.
17. Irish, J.C. et al., Amplification of WHSC1L1 regulates expression and estrogen-independent activation of ERα in SUM-44 breast cancer cells and is associated with ERα over-expression in breast cancer. Mol Oncol. 2016 Jun; 10(6): 850–865.
18. Hardiman G. et al., Systems analysis of the prostate transcriptome in African-American men compared with European-American men. Pharmacogenomics. 2016 Jul;17(10):1129-1143.
19. Allen, C., Kuhn, B.N., Cannella, N., Crow, A.D., Roberts, A.T., Lunerti, V., Ubaldi, M., Hardiman, G., Solberg Woods, L.C., Ciccocioppo, R., Kalivas, P.W., Chung, D., 2021.

Network-Based Discovery of Opioid Use Vulnerability in Rats Using the Bayesian Stochastic Block Model. Front Psychiatry 12, 745468.
